# Characterization of adult hippocampal neurogenesis in the novel *App^SAA^* Knock-in Alzheimer’s disease mouse model

**DOI:** 10.1101/2025.09.18.677216

**Authors:** Thomas A. Kim, Michelle D. Syty, Faye Wang, Shaoyu Ge

## Abstract

Adult hippocampal neurogenesis (AHN) declines with age and is thought to be severely exacerbated in neurodegenerative disorders like Alzheimer’s disease (AD). Despite numerous efforts to understand how AHN is altered in AD mouse models, results have been inconsistent, largely due to limitations of first-generation transgenic AD mouse models. The newly developed *App* Knock-in models address many of these limitations. Here, we provide the first in-depth characterization of hippocampal cell populations and AHN across different ages in the novel *App*^SAA^Knock-in (AppKI) mouse model. Our findings reveal that AppKI mice show no early deficits in excitatory dentate granule cells or inhibitory interneurons at 2 and 4 months, but significant loss of both populations emerges by 6 months of age. We also identified a progressive decline in AHN with the survival of newborn neurons being impaired first at 4 months, followed by a deficit in proliferation at 6 months. Furthermore, we demonstrate that exposure to enriched environment, a form of hippocampus-engaged exploration, robustly enhances AHN in AppKI mice, primarily by promoting survival. In conclusion, our study provides a foundational characterization of the AppKI model, establishing a timeline for cellular and AHN deficits.

## Introduction

The adult mammalian brain retains the capacity to generate new, functional neurons in the dentate gyrus (DG) of the hippocampus, a process termed adult hippocampal neurogenesis (AHN) (Eriksson et al., 1998; Kim et al., 2022; Toda et al., 2019). These newborn dentate granule cells (DGCs) integrate into existing neural circuits and are critical for hippocampal-dependent functions, including learning, memory, and pattern separation (Eriksson et al., 1998; Kim et al., 2022; Yang et al., 2020). Numerous studies have shown that the rate of AHN gradually decreases with age, a decline further accelerated in Alzheimer’s disease (AD) (Babcock et al., 2021; Moreno-Jiménez et al., 2019). However, despite extensive research, a clear consensus on how AD pathology affects AHN throughout aging remains elusive (Chuang, 2010; Kim et al., 2022).

Much of the existing literature provides inconsistent and even contradictory findings, with reports of increased, decreased, or unchanged neurogenesis in various AD mouse models (Kim et al., 2022; Zhou et al., 2023). These discrepancies can be largely attributed to the limitation of first-generation transgenic AD mouse models, which rely on the overexpression of mutant human proteins like *APP* or *Presenilin 1* (PSEN1) to mirror AD pathology in humans (Götz et al., 2018; Kim et al., 2022). Such models can produce non-physiological artifacts and confounding factors such as hyperactivity and seizure activity, which can affect the neural stem cell pool, complicating interpretation (Fu et al., 2019; Kim et al., 2022; Wang et al., 2022; Webster et al., 2014).

To overcome these challenges, second-generation knock-in (KI) models have been developed. In particular, the novel *App*SAA knock-in (AppKI) model introduces a humanized amyloid beta (Aβ) sequence with three familial AD mutations (Swedish, Arctic, and Austrian) (Xia et al., 2022). This ensures that mutant APP is expressed at physiological levels under its native promoter, providing a more accurate recapitulation of the human preclinical condition (Saito et al., 2014; Xia et al., 2022). This AppKI model presents with Aβ starting at 4 months of age (Kim et al., 2025; Xia et al., 2022), followed by robust neuroinflammation at 8 months and delayed behavioral and cognitive decline (Lu et al., 2025; Xia et al., 2022). This extended preclinical window makes it an ideal system for studying the early cellular events that precede cognitive decline.

Within the hippocampal neurogenic niche, the local circuitry, including inhibitory interneurons, plays a crucial role in regulating the proliferation, survival, and maturation of adult-born cells (Salta et al., 2023; Song et al., 2013). These interneurons are known to be severely affected in AD (Palop et al., 2007; Palop & Mucke, 2016; Verret et al., 2012), and recent work in knock-in models has identified them as a major contributor to amyloid plaque pathology (Rice et al., 2020). Therefore, a comprehensive analysis of AD’s impact on AHN must also consider the integrity of the surrounding cellular environment.

This study aimed to provide the first systematic characterization of AHN in the AppKI mouse model across different ages. Using immunofluorescence staining and BrdU labeling, we compared the integrity of the DG cell populations and the distinct stages of neurogenesis between wild-type (WT) and AppKI mice. We found that AHN is impaired as early as 4 months of age, coinciding with the initial formation of Aβ in this model. We further elucidate that the survival of newborn DGCs is affected first, followed by a decline in proliferation. Finally, given that non-pharmacological interventions like an enriched environment (EE) can enhance AHN (Nilsson et al., 1999; Shen et al., 2019; Wang et al., 2020), we investigated whether EE could rescue the AHN deficits in AppKI mice. We found that exposure to EE successfully increases AHN, primarily by promoting the survival of newborn neurons.

## Results

### Progressive loss of DGCs and inhibitory interneurons in AppKI mice begins at 6 months of age

The dentate gyrus (DG) is a subregion of the hippocampus harboring a diverse population of cell types, including excitatory dentate granule cells (DGCs) and inhibitory GABAergic interneurons (Amaral et al., 2007). Alzheimer’s disease (AD) patients exhibit progressive neuronal loss, and the hippocampus is particularly affected (Niikura et al., 2006; Rao et al., 2022). Similarly, many AD mouse models display neuronal loss as a common characteristic (Kim et al., 2022; Wirths & Bayer, 2010). To investigate this in the *App*SAA knock-in (AppKI) (Xia et al., 2022), we quantified the number of DGCs and interneurons in the DG of wild-type (WT) and AppKI mice.

First, we immunostained for prospero homeobox protein 1 (PROX1), a marker for immature and mature DGCs in the granule cell layer (GCL). We quantified the number of PROX1^+^ cells within a defined field of view and measured the thickness (dorsal-ventral axis) of the suprapyramidal blade of the GCL in 2-, 4-, 6-, and 9-month-old mice. At 2 and 4 months of age, no significant differences were observed between WT and AppKI mice in either the number of PROX1^+^ cells or the thickness of the GCL (**Supple. Fig. 1A-C; Fig. 1A-C**). However, by 6 months of age, AppKI mice exhibited a decrease in both parameters compared to WT mice (**Fig. 1D-F**). At 9 months, AppKI mice showed a markedly thinner GCL, while the number of PROX1^+^ cells displayed a downward trend (**Supple. Fig. 1D-F**). Notably, WT mice displayed a consistent age-related increase in both PROX1^+^ cells and GCL thickness (**Supple. Fig. 1G,H**), reflecting the continuous addition of new neurons. This developmental expansion was absent in AppKI mice (**Supple. Fig. 1I,J**).

**Figure 1.**
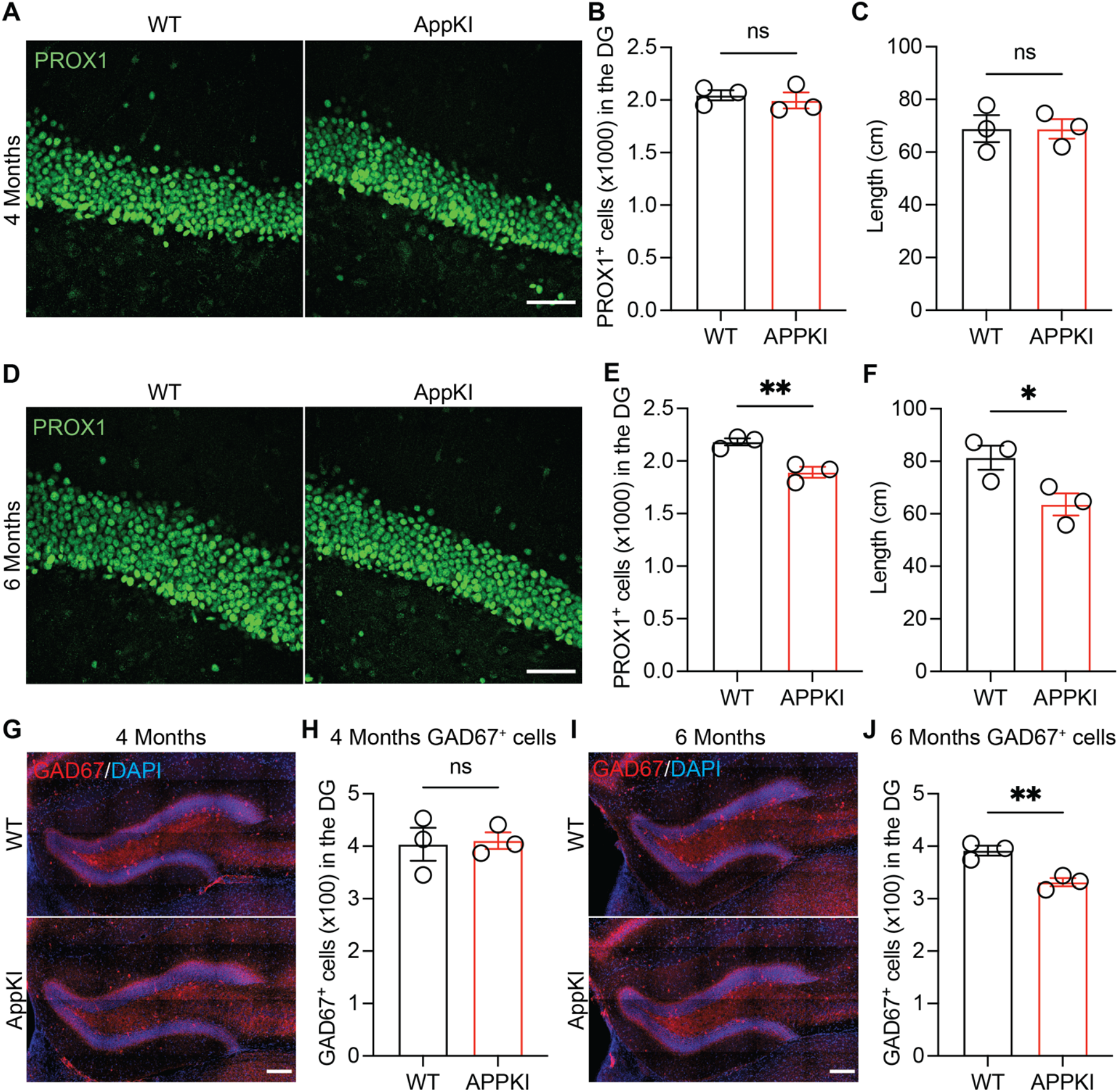
Analysis of PROX1^+^ cells and GAD67^+^ cells in the DG of 4- and 6-month-old WT and AppKI mice. **A** Representative confocal images of PROX1 staining of 4-month-old WT and AppKI mice in the DG (scale bar: 50 µm). **B** Quantification of PROX1^+^ cells of 4-month-old WT and AppKI mice in the suprapyramidal blade of the GCL (WT = 3, AppKI = 3; 2-tailed unpaired t-test; ns, p>0.05). **C** Analysis of the thickness (dorsal-ventral axis) of the suprapyramidal blade of the PROX1^+^ cells in 4-month-old WT and AppKI mice (WT = 3, AppKI = 3; 2-tailed unpaired t-test; ns, p>0.05). **D** Representative confocal images of PROX1 staining of 6-month-old WT and AppKI mice in the DG (scale bar: 200 µm). **E** Quantification of PROX1^+^ cells of 6-month-old WT and AppKI mice in the suprapyramidal blade of the GCL (WT = 3, AppKI = 3; 2-tailed unpaired t-test; **p<0.01). **F** Analysis of the thickness (dorsal-ventral axis) of the suprapyramidal blade of the PROX1^+^ cells in 6-month-old WT and AppKI mice (WT = 3, AppKI = 3; 2-tailed unpaired t-test; *p<0.05). **G** Representative confocal images of GAD67 staining of 4-month-old WT and AppKI mice in the DG (scale bar: 200 µm). **H** Quantification of GAD67^+^ cells of 4-month-old WT and AppKI mice in the DG (WT = 3, AppKI = 3; 2-tailed unpaired t-test; ns, p>0.05). **I** Representative images of GAD67 staining of 6-month-old WT and AppKI mice in the DG (scale bar: 200 µm). **J** Quantification of GAD67^+^ cells of 6-month-old WT and AppKI mice in the DG (WT = 3, AppKI = 3; 2-tailed unpaired t-test; **p<0.01).

Next, we investigated GABAergic interneurons by immunostaining hippocampal slices for glutamic acid decarboxylase 67 (GAD67). We quantified the number of GAD67^+^ cells within the DG of 2-, 4-, 6-, and 9-month-old mice. No significant differences were observed between WT and AppKI mice at 2 and 4 months of age (**Supple. Fig. 2A,B; Figure 1G,H**). However, a significant decrease in GAD67^+^ cells was evident in AppKI mice compared to WT mice at 6 months of age (**Fig. 1I,J**). Interestingly, this difference was no longer significant by 9 months of age, as WT mice began to show an age-related decline (**Supple. Fig. 2C,D**). This suggests an accelerated loss of GAD67+ interneurons in AppKI mice, with a decline beginning around 6 months, whereas in WT mice, this loss starts later around 9 months (**Supple. Fig. 2E,F**).

Put together, our findings indicate that the AppKI model does not exhibit early developmental deficits in excitatory or inhibitory neuron populations. However, beginning at 6 months, AppKI mice show a clear deficit in DGCs and an accelerated loss of inhibitory interneurons compared to WT mice.

### Adult hippocampal neurogenesis is impaired in AppKI mice starting at 4 months of age

The failure of the GCL to expand in AppKI mice, coupled with the loss of interneurons known to regulate the neurogenic niche (Masiulis et al., 2011; Salta et al., 2023), strongly suggested a potential deficit in AHN. To investigate this directly, we characterized the population of newborn neurons across the same age range.

We immunostained hippocampal sections from 2-, 4-, 6-, and 9-month-old WT and AppKI mice with doublecortin (DCX), a marker for newborn neurons (**Fig. 2A**). We quantified DCX^+^ cells in the subgranular zone (SGZ) to estimate their total number in the DG. At 2 months, both genotypes showed robust neurogenesis with no significant difference in the number of DCX^+^ cells (**Fig. 2B**). As expected, the number of DCX^+^ cells declined sharply with age in both WT and AppKI mice (**Supple. Fig. 3A,B**). However, beginning at 4 months of age, AppKI mice had significantly fewer DCX^+^ cells compared to their age-matched WT counterparts (**Fig. 2C**).

**Figure 2.**
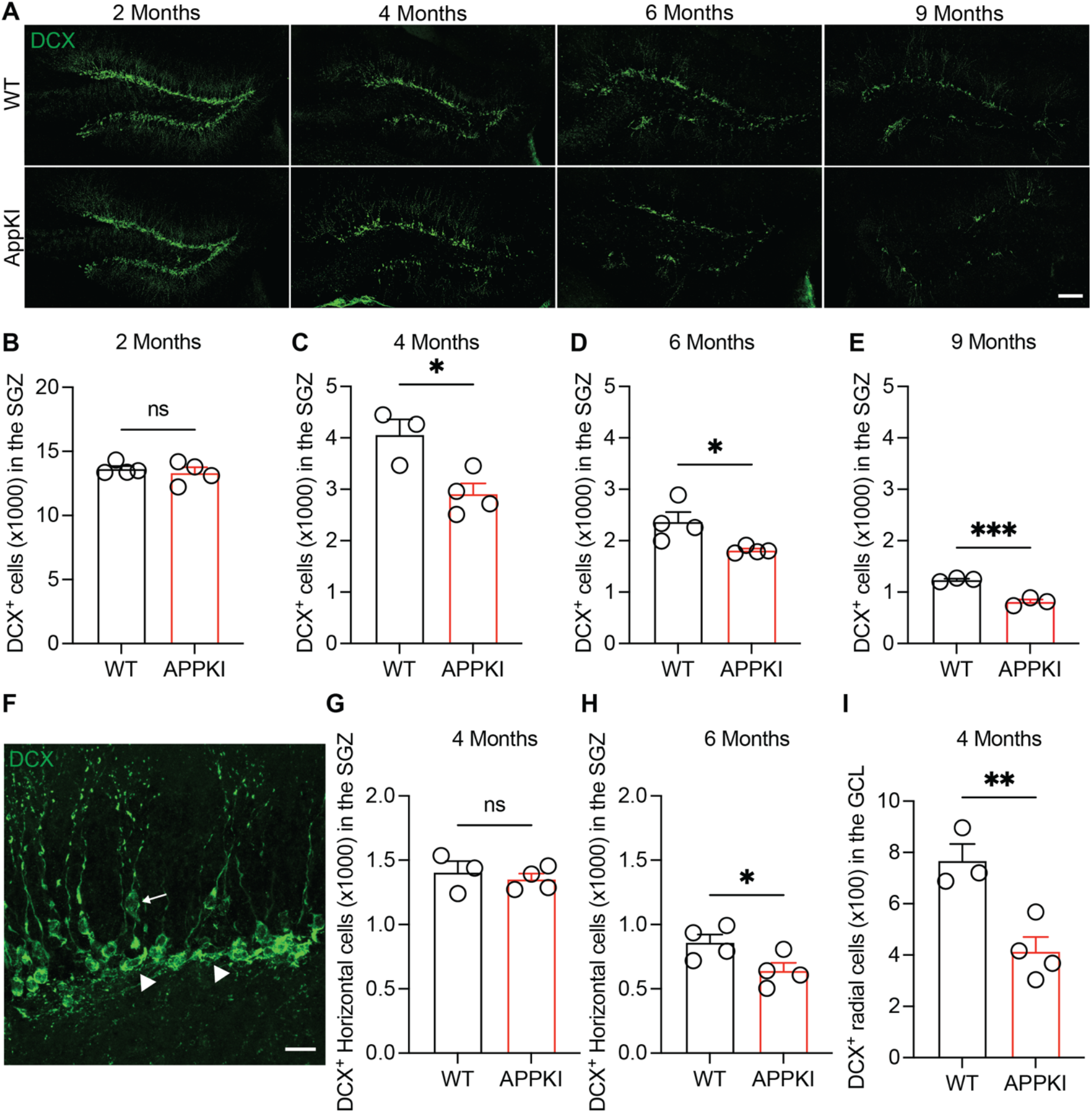
Impaired adult hippocampal neurogenesis in AppKI mice across age. **A** Representative confocal images of DCX immunostaining of 2-, 4-, 6-, and 9-month-old WT and AppKI mice in the DG (scale bar: 200 µm) **B** Quantification of DCX^+^ cells in the SGZ of 2-month-old WT and AppKI mice (WT =4, AppKI = 4; 2-tailed unpaired t-test; ns, p>0.05). **C** Quantification of DCX^+^ cells in the SGZ of 4-month-old WT and AppKI mice (WT = 3, AppKI = 4; 2-tailed unpaired t-test; *p<0.05). **D** Quantification of DCX^+^ cells in the SGZ of 6-month-old WT and AppKI mice (WT = 4, AppKI = 4; 2-tailed unpaired t-test; *p<0.05). **E** Quantification of DCX^+^ cells in the SGZ of 9-month-old WT and AppKI mice (WT = 3, AppKI = 3; 2-tailed unpaired t-test; ***p<0.001). **F** Representative confocal images of DCX^+^ horizontal cells (white arrowhead) and radial cells (white arrow) in the DG (scale bar: 20 µm). **G** Quantification of DCX^+^ horizontal cells in the SGZ of 4-month-old WT and AppKI mice (WT = 3, AppKI = 4; 2-tailed unpaired t-test; ns, p>0.05). **H** Quantification of DCX^+^ horizontal cells in the SGZ of 6-month-old WT and AppKI mice (WT = 4, AppKI = 4; 2-tailed unpaired t-test; *p<0.05). **I** Quantification of DCX^+^ radial cells in the GCL of 4-month-old WT and AppKI mice (WT = 3, AppKI = 4; 2-tailed unpaired t-test; **p<0.01).

This decrease persisted and remained significant at both 6 and 9 months of age. (**Fig. 2D,E**) To determine which specific stages of neuronal development were affected, we categorized DCX^+^ cells based on their morphology and location (Ge et al., 2006; Sun et al., 2015; Wang et al., 2019; Yang et al., 2020). Horizontal DCX^+^ cells in the SGZ represent an earlier, proliferative stage, while radial DCX^+^ cells extending into the GCL represent a later, survival stage (**Fig. 2F**). At 4 months, the number of horizontal DCX^+^ cells was comparable between genotypes (**Fig. 2G**). However, by 6 months, AppKI mice had significantly fewer horizontal DCX^+^ cells, suggesting an impairment in the proliferation of newborn DGCs (**Fig. 2H**). Notably, at 4 months, AppKI mice showed a significant reduction in the number of radial DCX^+^ cells in the GCL (**Fig. 2I**), indicating deficits in newborn DGC survival. This deficit was also present at 6 months (**Supple. Fig. 3C**).

Our findings collectively demonstrate a rapid decline in AHN in AppKI mice beginning at 4 months of age. This is further supported by the decrease in radial DCX^+^ cells within the GCL at 4 months in AppKI mice, suggesting a deficit in the survival of newborn DGCs. Additionally, the difference in the number of DCX^+^ horizontal cells observed at 6 months in AppKI mice, specifically suggests impaired proliferation.

### A sequential decline in newborn DGC survival and proliferation in AppKI mice

To confirm the sequential impairment suggested by the DCX data, we directly assessed the proliferation and survival rates of newborn DGCs in 4- and 6-month-old WT and AppKI mice using 5-bromo-2’-deoxyuridine (BrdU) labeling.

To assess proliferation, mice received a single intraperitoneal injection of BrdU (10ug/g) and were analyzed 48 hours later (**Fig. 3A**). At 4 months of age, the number of BrdU^+^ cells in the SGZ were comparable between WT and AppKI mice (**Supple. Fig. 4A**). However, by 6 months, AppKI mice showed a significant reduction in BrdU^+^ cells compared to WT controls (**Fig. 3B**). We confirmed this finding using endogenous markers of proliferation, MCM2 and TBR2 (Simard et al., 2025; Zhang & Jiao, 2015). Consistent with the BrdU data, the numbers of MCM2^+^ and TBR2^+^ cells were comparable between genotypes at 4 months (**Supple. Fig. 4B,C**) but were significantly decreased in AppKI mice at 6 months (**Fig. 3C,D**).

**Figure 3.**
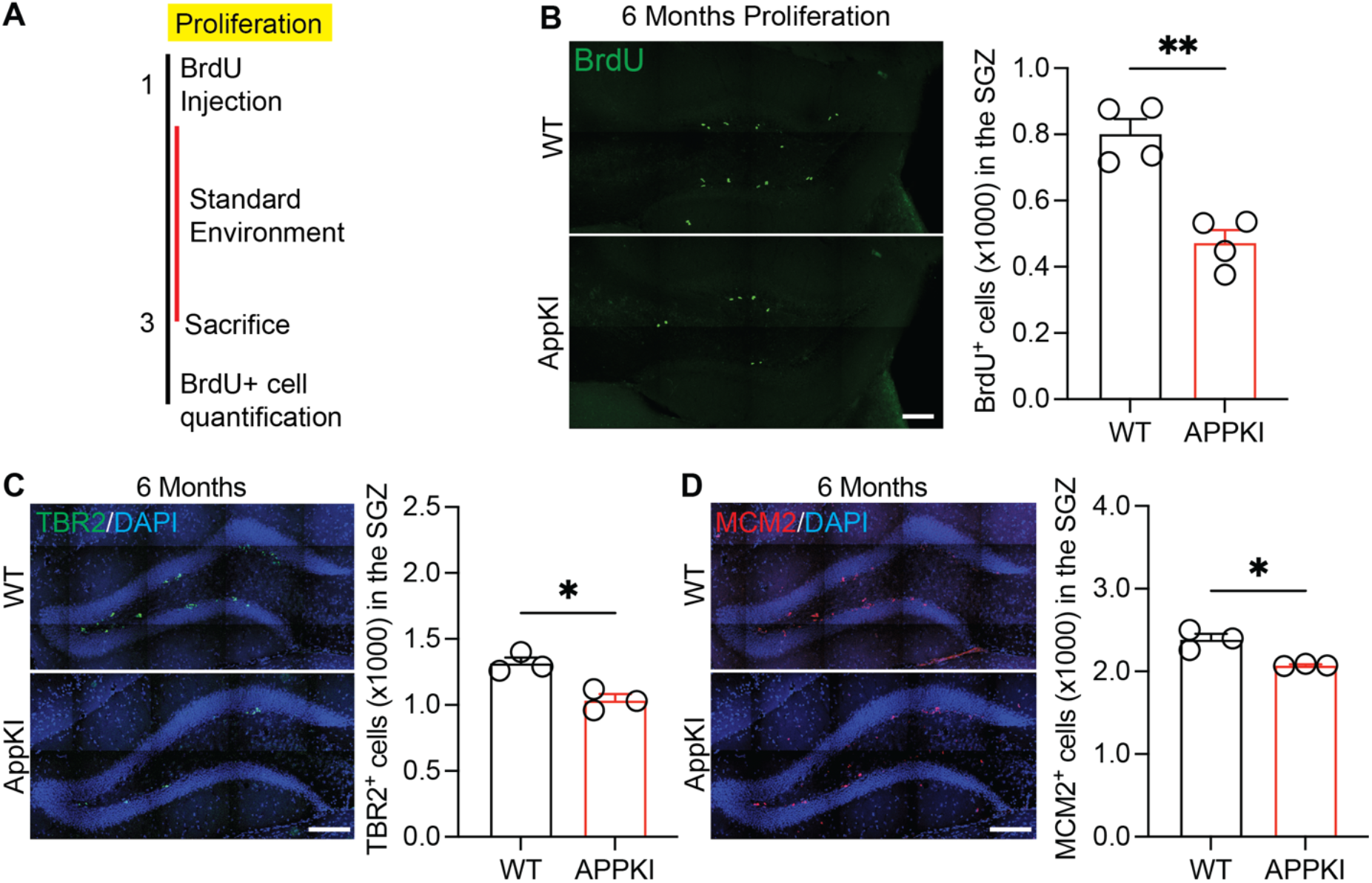
Declining proliferation of newborn neurons in 6-month-old WT and AppKI mice. **A** Schematic of the experimental paradigm for BrdU injections to examine the proliferation rate. **B Left:** Representative confocal images of BrdU immunostaining in the DG of 6-month-old WT and AppKI mice for proliferation rate analysis (scale bar: 100 µm). **Right:** Quantification of BrdU+ cells in the SGZ for proliferation rate of AHN in 6-month-old WT and AppKI mice (WT = 4, AppKI = 4; 2-tailed unpaired t-test; *p<0.01). **C Left:** Representative confocal images of TBR2 immunostaining in the DG of 6-month-old WT and AppKI mice (scale bar: 200 µm). **Right:** Quantification of TBR2^+^ cells in the SGZ of 6-month-old WT and AppKI mice (WT = 3, AppKI = 3; 2-tailed unpaired t-test; *p<0.05). **D Left:** Representative confocal images of MCM2 immunostaining in the DG of 6-month-old WT and AppKI mice (scale bar: 200 µm). **Right:** Quantification of MCM2^+^ cells in the SGZ of 6-month-old WT and AppKI mice (WT = 3, AppKI = 3; 2-tailed unpaired t-test; *p<0.05).

To assess survival, mice received a single BrdU injection and were analyzed 2 weeks later (**Fig. 4A**). BrdU^+^ cells in the SGZ were then quantified. Consistent with our DCX data, AppKI mice displayed a significantly lower number of surviving BrdU^+^ cells at both 4 and 6 months of age compared to WT mice (**Fig. 4B,C**).

**Figure 4.**
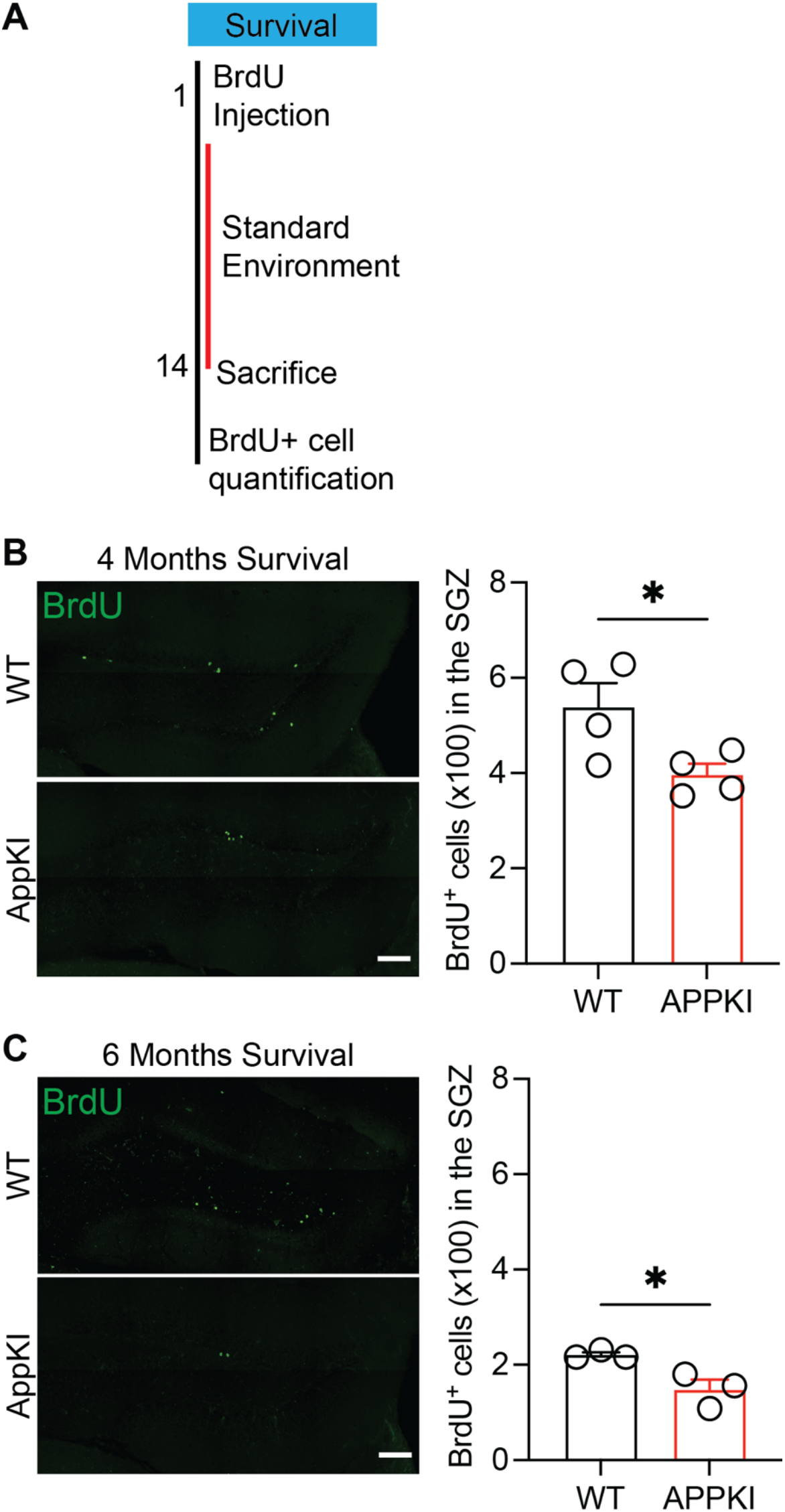
Declining survival of newborn neurons in 4- and 6-month-old WT and AppKI mice. **A** Schematic of the experimental paradigm for BrdU injections to examine the survival rate of newborn DGCs. **B Left:** Representative confocal images of BrdU immunostaining in the DG of 4-month-old WT and AppKI mice for survival rate analysis (scale bar: 200 µm). **Right:** Quantification of BrdU^+^ cells in the SGZ for survival rate of AHN in 4-month-old WT and AppKI mice (WT = 4, AppKI = 4; 2-tailed unpaired t-test; *p<0.05). **C Left:** Representative confocal images of BrdU immunostaining in the DG of 6-month-old WT and AppKI mice for survival rate analysis (scale bar: 200 µm). **Right:** Quantification of BrdU^+^ cells in the SGZ for survival rate of AHN in 6-month-old WT and AppKI mice (WT = 3, AppKI = 3; 2-tailed unpaired t-test; *p<0.05).

Taken together, these data confirm a sequential disruption of neurogenesis in the AppKI model. The survival of newborn DGCs is the first process to be impaired, with deficits appearing as early as 4 months. This is followed by a subsequent decline in proliferation of newborn neurons, which becomes evident at 6 months.

### Enriched environment increases adult hippocampal neurogenesis in 4- and 6-month-old AppKI mice

Given the clear impairment of AHN in AppKI mice, we next asked whether this deficit could be rescued by a non-pharmacological intervention. While enriched environments (EE) are known to enhance AHN in WT mice (Nilsson et al., 1999; Shen et al., 2019; Wang et al., 2020), their impact and efficacy in AD models remains controversial (Babcock et al., 2021; Kim et al., 2022). Therefore, we tested the effects of a short-term EE exposure at two critical time points: 4 months, when the AHN decline begins (**Fig. 2C**), and 6 months, when the deficit is well-established (**Fig. 2D**).

4- and 6-month-old AppKI mice were housed in either standard (STD) cages with bedding, food, and water or EE cages equipped with 5-6 additional objects for 3 days. Subsequently, the mice were sacrificed, and their hippocampal slices were immunostained for DCX. At both 4 and 6 months of age, AppKI mice housed in EE cages showed a significant increase in the total number of DCX^+^ cells compared to their counterparts in standard housing (**Fig. 5A-C**). To determine which stage of neurogenesis was affected by this intervention, we again analyzed the horizontal and radial DCX^+^ cell populations. In both 4- and 6-month-old AppKI mice, EE had no significant effect on the number of horizontal DCX^+^ cells (**Fig. 5D,E**). However, EE treatment led to a significant increase in the number of radial DCX^+^ cells in the GCL at both ages (**Fig. 5F,G**).

**Figure 5.**
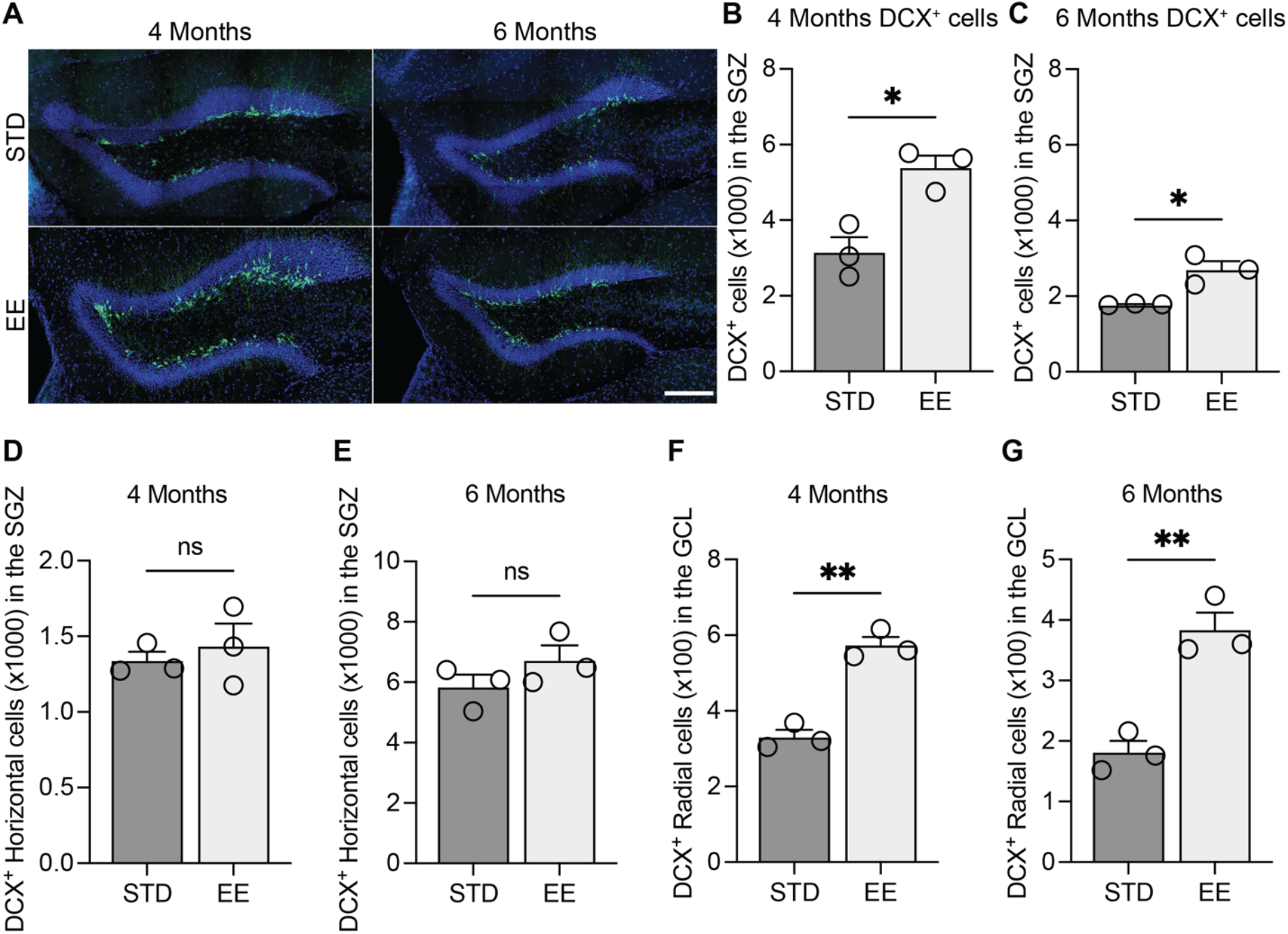
Enriched environment enhances AHN in the SGZ of 4- and 6-month-old AppKI mice. **A** Representative confocal images of DCX immunostaining in the SGZ of 4- and 6-month-old AppKI mice in STD and EE settings (scale bar: 200um). **B** Quantification of DCX^+^ cells in the SGZ zone of 4-month-old AppKI mice in STD and EE settings (STD = 3, EE = 3; 2-tailed unpaired t-test; *p<0.05). **C** Quantification of DCX^+^ cells in the SGZ zone of 6-month-old AppKI mice in STD and EE settings (STD = 3, EE = 3; 2-tailed unpaired t-test; *p<0.05). **D** Quantification of DCX^+^ horizontal cells in the SGZ of 4-month-old AppKI mice in STD and EE settings (STD = 3, EE = 3; 2-tailed unpaired t-test; ns, p>0.05). **E** Quantification of DCX^+^ horizontal cells in the SGZ of 6-month-old AppKI mice in STD and EE settings (STD = 3, EE = 3; 2-tailed unpaired t-test; ns, p>0.05). **F** Quantification of DCX^+^ radial cells in the GCL of 4-month-old AppKI mice in STD and EE settings (STD = 3, EE = 3; 2-tailed unpaired t-test; **p<0.01). **G** Quantification of DCX^+^ radial cells in the GCL of 6-month-old AppKI mice in STD and EE settings (STD = 3, EE = 3; 2-tailed unpaired t-test; **p<0.01).

While previous studies investigating the effects of EE on AHN in AD models have yielded mixed results, our findings provide compelling evidence for the beneficial impact of EE on AHN in the AppKI mouse model. Moreover, our analyses on horizontal and radial DCX^+^ cells suggest that EE specifically impacts the survival of newborn neurons rather than the proliferation. This finding adds valuable nuance to the current understanding of EE’s role in AD models, suggesting its potential therapeutic benefit at specific stages of disease progression and highlighting the importance of considering model-specific characteristics.

## Discussion

In this study, we provide the first systematic characterization of adult hippocampal neurogenesis in the *App*^SAA^Knock-in (AppKI) mouse model of AD. Our results establish a clear, age-dependent cellular deficit in the dentate gyrus with accelerated neuronal loss of both excitatory DGCs and GABAergic inhibitory interneurons in the AppKI mouse model. We also identify a sequential impairment of neurogenesis in the AppKI mice starting at 4 months of age, evidenced by fewer DCX^+^ cells in the SGZ. With further analysis of the proliferation and survival rate of newborn DGCs, we discovered that there was an initial reduction in the survival rate of DGCs at 4 months of age, followed by a decline in both proliferation and survival of DGCs at 6 months of age in the AppKI mice. Finally, we demonstrate that EE increases DCX^+^ cells in both 4- and 6-month-old AppKI mice. These findings offer crucial insights into the early pathophysiology of AD and highlights the therapeutic potential for targeting neurogenesis in AD.

A key contribution of this work is the foundational characterization of the novel AppKI model, which avoids the artifacts of *APP* overexpression (Saito et al., 2014; Xia et al., 2022). Here, we establish a physiologically relevant timeline of neurodegeneration by using immunostaining to quantify and analyze the changes in excitatory DGCs and GABAergic inhibitory interneurons in the AppKI mouse model across age. We found no evidence of early developmental deficits, with DGC and interneuron numbers being comparable between genotypes at 2 and 4 months. The onset of significant neuronal loss at 6 months occurs after the initial appearance of Aβ at 4 months (Kim et al., 2025; Xia et al., 2022) and precedes the established cognitive decline at 12 months in this model (Lu et al., 2025). This positions cellular loss as an intermediate event, bridging the gap between initial amyloid pathology and eventual cognitive impairment. However, several limitations should be considered. First, our study relies solely on cell number quantification via immunostaining. Future experiments with techniques such as electrophysiology or functional imaging could provide deeper insights into neuronal function and potential synaptic deficits (Gu et al., 2011). Furthermore, the study focused on the cell populations, overlooking variations in specific interneuron subtypes that may show unique vulnerabilities. Delving deeper into specific interneuron subtypes using targeted markers could reveal their individual roles and susceptibilities linking them to potential therapeutic interventions. In addition, the ages examined were not continuous which could lead to missing important transient changes or the full progression of neuronal loss. Employing longitudinal studies that track neuronal populations over time would capture the dynamic changes and pinpoint critical transition points in the disease process.

Our most striking finding is the sequential impairment of AHN. We found a significant decrease in DCX^+^ cells in the SGZ of AppKI mice starting at 4 months of age. Importantly, we observed differences in horizontal and radial DCX^+^ cell populations in AppKI mice. 6-month-old AppKI mice had fewer horizontal DCX^+^ cells, suggesting impairment in intermediate progenitor cells or the proliferation process. On the other hand, 4-month-old AppKI exhibited a noticeably reduced number of radial DCX^+^ cells in the GCL, hinting at potential deficits in newborn DGCs migration. These results were supported with BrdU, MCM2, and TBR2 analyses. This specific timeline helps resolve some of the inconsistencies in the literature, which often reports on a single time point or a single stage of neurogenesis. Nevertheless, this study focuses on the descriptive aspects of AHN. Further studies exploring the precise mechanisms behind these changes in proliferation and survival of newborn DGCs are crucial to understanding the early impairment of AHN in AppKI mice. Employing additional approaches, such as live imaging, could offer valuable insights into the dynamics of neurogenesis. Additionally, exploring specific signaling pathways involved in neurogenesis might shed light on the key regulatory factors driving its impairment in AD. Furthermore, utilizing retroviral labeling of newborn DGCs would offer more comprehensive information on proliferation, migration, and integration, remaining as a crucial experiment to perform in the near future (Gu et al., 2012).

EE is known to play a crucial role in AHN and improve performance in behavioral tasks in WT mice (Nilsson et al., 1999; Shen et al., 2019; Wang et al., 2020). However, its efficacy in AD models remains debated due to inconsistent results (Babcock et al., 2021; Kim et al., 2022). Our findings stand out, demonstrating that EE significantly increases DCX^+^ cells in both 4- and 6-month-old AppKI mice, providing compelling evidence for its potential benefits in AD models. Moreover, we observed that EE impacts the survival of newborn neurons in the GCL significantly. However, further research is needed to fully understand the long-term impact of EE and its relationship with cognitive function. Future studies should investigate the impact of different exposure durations to identify an optimal timeframe for beneficial effects. Additionally, exploring the correlation between EE-mediated neurogenesis and cognitive function through behavioral assessments will be critical.

Despite these limitations, our study establishes the AppKI mouse as a valuable model for studying AHN in AD. We define a precise timeline for cellular pathology and AHN decline, demonstrating that these are early features of the disease, and show that this deficit is amenable to rescue by environmental stimulation. These findings underscore the importance of AHN as a contributor to AD pathogenesis and as a promising target for therapeutic interventions.

## Materials and Methods

### Mice

All surgeries and experimental procedures were performed as approved by the Stony Brook University Animal Use Committee and in accordance with the guidelines of the National Institutes of Health. Experiments were conducted using male and female 2, 4, 6, and 9 months C57BL/6J wildtype mice (Jackson Laboratory, Stock No. 000664) and *App*^SAA^KI mice on a C57BL/6J background (Jackson Laboratory, Stock No. 034711). All mice were maintained on a 12-hour light/dark cycle with *ad libitum* access to food and water.

### Perfusion and Tissue processing

All mice were deeply anesthetized with urethane (200ug/g) and transcardially perfused with 0.1 M PBS and then 4% PFA. Brains were removed, fixed overnight in 4% PFA and then transferred to a 30% (w/v) sucrose solution and stored at 4°C until sectioning. Brains were sectioned into 40 μm coronal sections covering the entire anterior/posterior axis of the DG. Sections were stored in cryopreservative until used for immunostaining.

### Immunohistochemistry

For all BrdU labeling experiments, sections were pretreated with 2N HCL for 25 minutes at 37°C followed by 2x 10 minutes in 0.1 M sodium borate (pH = 8.5) at room temperature. For all MCM2 stainings, sections were pretreated with 0.1% citrate buffer solution for heat-induced antigen retrieval. Sections were first exposed to 0.5% Triton X-100 in 0.1M PBS (PBST) for 10 min for the permeabilization step. We then blocked the sections with 10% donkey serum in 0.25% PBST for 1 h at room temperature. The sections were incubated overnight at 4°C with primary antibody in 0.25% PBST and 3% donkey serum. The primary antibodies used in this dissertation include BrdU (rat monoclonal antibody, 1:500, Abcam, ab6326), Doublecortin (DCX, rabbit polycolonal antibody, 1:1000, Cell Signaling, 4604), GAD67 (Mouse monoclonal antibody, 1:250, Millipore-sigma, MAB5406), MCM2 (mouse monoclonal antibody, 1:500, BD Biosciences, BD 610701), PROX1 (rabbit polyclonal antibody, 1:500 Millipore-sigma, AB5475), and TBR2/Eomes (rabbit polyclonal antibody, 1:250, Abcam, ab23345). The following day, the sections were washed 3x 10min in 0.25% PBST and incubated with secondary antibodies Alexa Fluor 488-conjugated donkey anti-rabbit (1:1000; Jackson Immuno Research, AB_2313584), Alexa, Fluor 488-conjugated donkey anti-rat (1:1000, Invitrogen, A21208), and Alexa Fluor 568-conjugated donkey anti-mouse (1:250, Invitrogen, A10037) in 0.25% PBST and 3% donkey serum for 3 h while shaking at room temperature. Sections were mounted with DAPI fluoromount, and images were obtained on a Zeiss LSM800 confocal microscope.

### 5-Bromo-2’ Deoxyuridine (BrdU) Administration

A single dosage of 10µl/g BrdU was administered intraperitoneally (i.p.) for the analysis of proliferation rate and survival rate of newborn DGCs. The mice were euthanized 48 hours later for proliferation experiments and two weeks later for survival experiments.

### Quantifications

#### PROX1 Quantification

For PROX1, we counted the number of fluorescent cells in the middle 1/3rd of the suprapyramidal blade of the GCL of every tenth brain section. We then multiplied this number by ten to arrive at an approximation for the total number of labeled cells in the suprapyramidal blade.

#### GAD67 Quantification

For GAD67, we counted the number of fluorescent cells in all three layers of the DG of every tenth brain section. We then multiplied this number by ten to arrive at an approximation for the total number of labeled cells.

#### DCX Quantification

For DCX density analysis, we selected every eighth brain section throughout the anterior/posterior axis of the DG, counted the number of DCX^+^ cells in the middle 1/3rd of the suprapyramidal and infrapyramidal blades. We then multiplied this number by eight to arrive at an approximation for the total number of labeled cells.

#### BrdU Quantification

For BrdU analyses, we used every fourth brain section and counted fluorescently labeled cells within the SGZ across the entire anterior-posterior axis of the DG. We then multiplied this number by four to arrive at an approximation for the total number of labeled cells in the SGZ.

#### TBR2 and MCM2 Quantification

For TBR2 and MCM2 analysis, we counted the number of fluorescent cells across the entire SGZ of every tenth brain section across the entire anterior-posterior axis of the DG. We then multiplied this number by ten to arrive at an approximation for the total number of labeled cells.

### Enriched environment exploration

4- and 6-month-old AppKI mice were housed in either a standard cage with bedding, food and water, or an EE cage which consisted of 5-6 novel objects for the mice to explore for three days.

## Acknowledgements

We thank all members of the Ge and Xiong laboratories for their valuable comments. This work was supported by R01AG066912 to S.G.

## Statistical Analysis

Mice were randomly assigned to experimental or control groups within litters. Each experiment will be repeated at least three times to ensure reproducibility. Sample sizes are based on preliminary results or prior studies to achieve >85% power for detecting significant differences. Two-tailed unpaired t-test was used for all statistical analyses. All statistical analyses were conducted with 95% confidence intervals, and p-values of 0.05 were considered as the cutoff for statistical significance. Statistical analysis was performed using Prism 10.0.3 (GraphPad Software, Boston, Massachusetts). All data are presented as mean ± standard error of the mean (SEM).

## Author Contributions

Conceptualization, T.A.K. and S. Ge; Methodology, T.A.K. and S. Ge; Formal Analysis T.A.K., M.D.S., and F.W.; Investigation, T.A.K., M.D.S., F.W.; Writing, T.A.K. and S. Ge produced the initial manuscript, and all other authors reviewed and commented on the manuscript; Supervision, S. Ge; Funding Acquisition, S.Ge.

