## Supplemental Figures for "Characterization of adult hippocampal neurogenesis in the novel *App^SAA^* Knock-in Alzheimer’s disease mouse model"

**Supplementary Figures**

#### Supplementary Figure 1

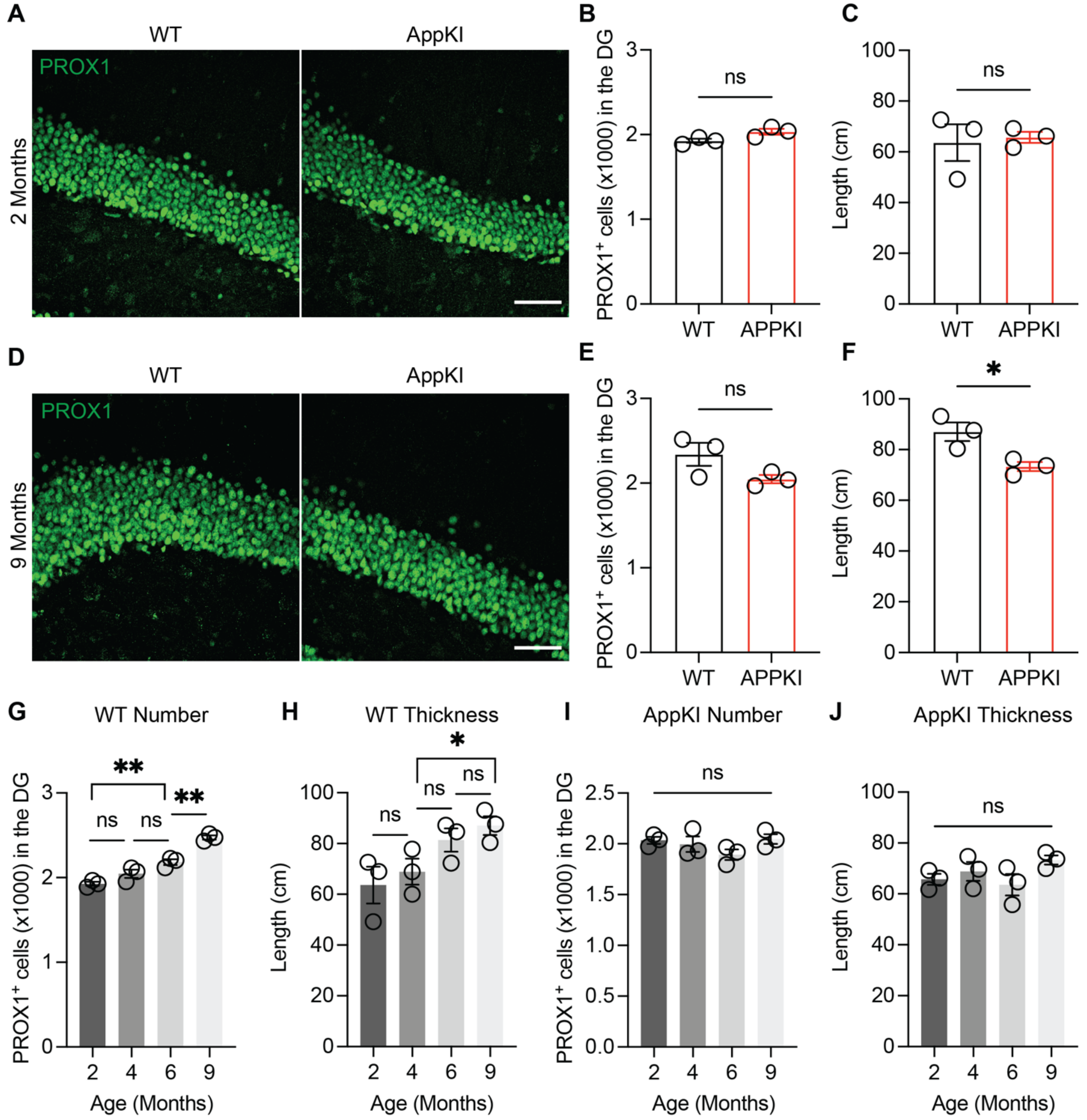


**Supplementary Figure 1 Characterization of PROX1$^{+}$ cells in the DG of 2- and 9-month-old WT and AppKI mice**

**A.** Representative confocal images of PROX1 staining of 2-month-old WT and AppKI mice in the DG (scale bar: 50 µm).

**B.** Quantification of PROX1$^+$ cells of 2-month-old WT and AppKI mice in the suprapyramidal blade of the GCL (WT = 3, AppKI = 3; 2-tailed unpaired t-test; ns, $p>0.05$).

**C**. Analysis of the thickness (dorsal-ventral axis) of the suprapyramidal blade of the GCL in 2-month-old WT and AppKI mice (WT = 3, AppKI = 3; 2-tailed unpaired t-test; ns, $p>0.05$).

**D.** Representative confocal images of PROX1 staining of 9-month-old WT and AppKI mice in the DG (scale bar: 50 µm).

**E.** Quantification of PROX1$^+$ cells of 9-month-old WT and AppKI mice in the suprapyramidal blade of the GCL (WT = 3, AppKI = 3; 2-tailed unpaired t-test; ns, $p>0.05$).

**F.** Analysis of the thickness (dorsal-ventral axis) of the suprapyramidal blade of the GCL in 9-month-old WT and AppKI mice (WT = 3, AppKI = 3; 2-tailed unpaired t-test; *$p<0.05$).

**G.** Analysis of PROX1$^+$ cell number in the suprapyramidal blade of the GCL in WT mice across age (n = 3 for all age groups; 2-tailed unpaired t-test; ns, $p>0.05$, **$p<0.01$).

**H.** Analysis of thickness (dorsal-ventral axis) of the suprapyramidal blade of the PROX1$^+$ cells across age in WT mice (n = 3 for all age groups; 2-tailed unpaired t-test; ns, $p>0.05$, *$p<0.05$).

**I.** Analysis of PROX1$^+$ cell number in the suprapyramidal blade of the GCL in AppKI mice across age (n = 3 for all age groups; 2-tailed unpaired t-test; ns, $p>0.05$).

**J.** Analysis of thickness (dorsal-ventral axis) of the suprapyramidal blade of the PROX1$^+$ cells across age in AppKI mice (n = 3 for all age groups; 2-tailed unpaired t-test; ns, $p>0.05$).

### Supplementary Figure 2

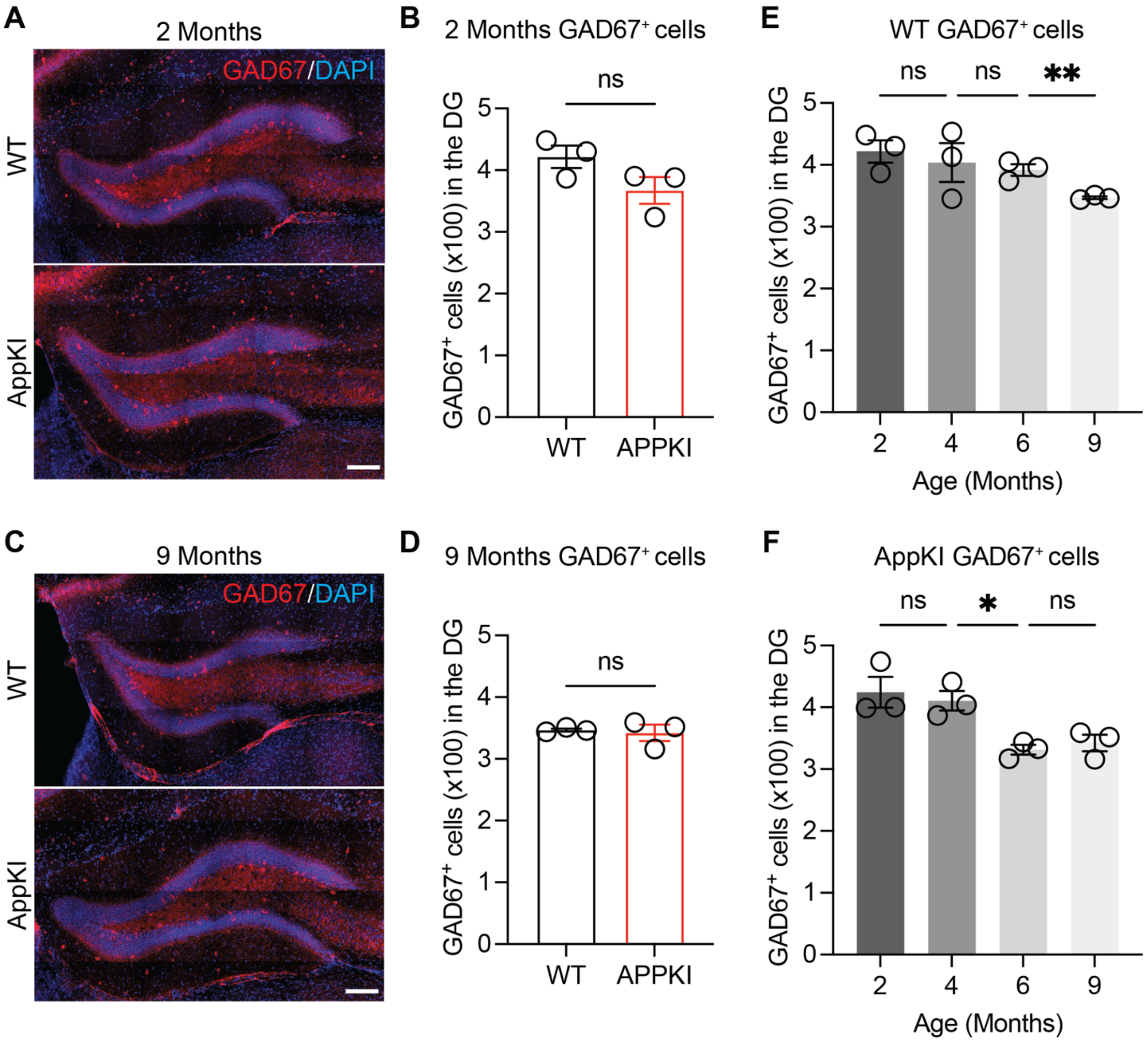


**Supplementary Figure 2 Characterization of GAD67$^{+}$ cells in the DG of 2- and 9-month-old WT and AppKI mice**

**A** Representative confocal images of GAD67 staining of 2-month-old WT and AppKI mice in the DG (scale bar: 200 μm).

**B** Quantification of GAD67$^{+}$ cells of 2-month-old WT and AppKI mice in the DG (WT = 3, AppKI = 3; 2-tailed unpaired t-test; ns, p>0.05).

**C** Representative images of GAD67 staining of 9-month-old WT and AppKI mice in the DG (scale bar: 200 μm)

**D** Quantification of $GAD67^+$ cells of 9-month-old WT and AppKI mice in the DG (WT = 3, AppKI = 3; 2-tailed unpaired t-test; ns, $p>0.05$).

**E** Quantification of $GAD67^+$ cells across age in WT mice (n = 3 for all age groups; 2-tailed unpaired t-test; ns, $p>0.05$, $^{**}p<0.01$).

**F** Quantification of $GAD67^+$ cells across age in AppKI mice (n = 3 for all age groups; 2-tailed unpaired t-test; ns, $p>0.05$, $^{*}p<0.05$).

### Supplementary Figure 3

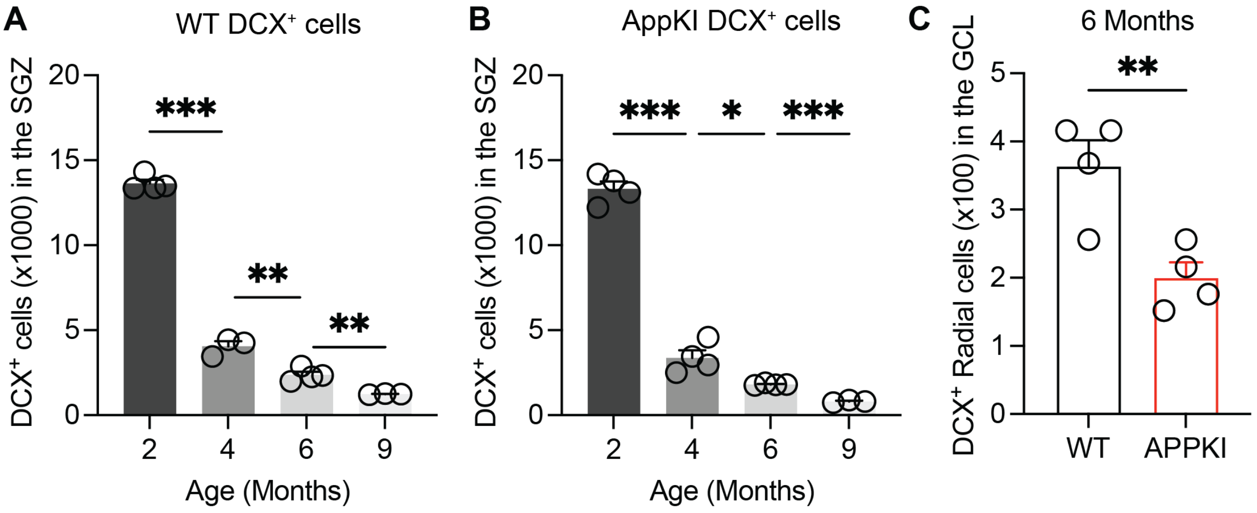


**Supplementary Figure 3 Characterization of DCX$^+$ cells in the DG of WT and AppKI mice across age**

**A** Quantification of DCX$^+$ cells in the SGZ of WT mice across age (2, 6 months = 4 mice, 4, 9 months = 3 mice; 2-tailed unpaired t-test; **$p<0.01$, ***$p<0.001$).

**B** Quantification of DCX$^+$ cells in the SGZ of AppKI mice across age (2, 4, 6 months = 4 mice, 9 months = 3 mice; 2-tailed unpaired t-test; *$p<0.05$, ***$p<0.001$).

**C** Quantification of DCX$^+$ radial cells in the GCL of 6-month-old WT and AppKI mice (WT = 4, AppKI = 4; 2-tailed unpaired t-test; **$p<0.01$).

### Supplementary Figure 4

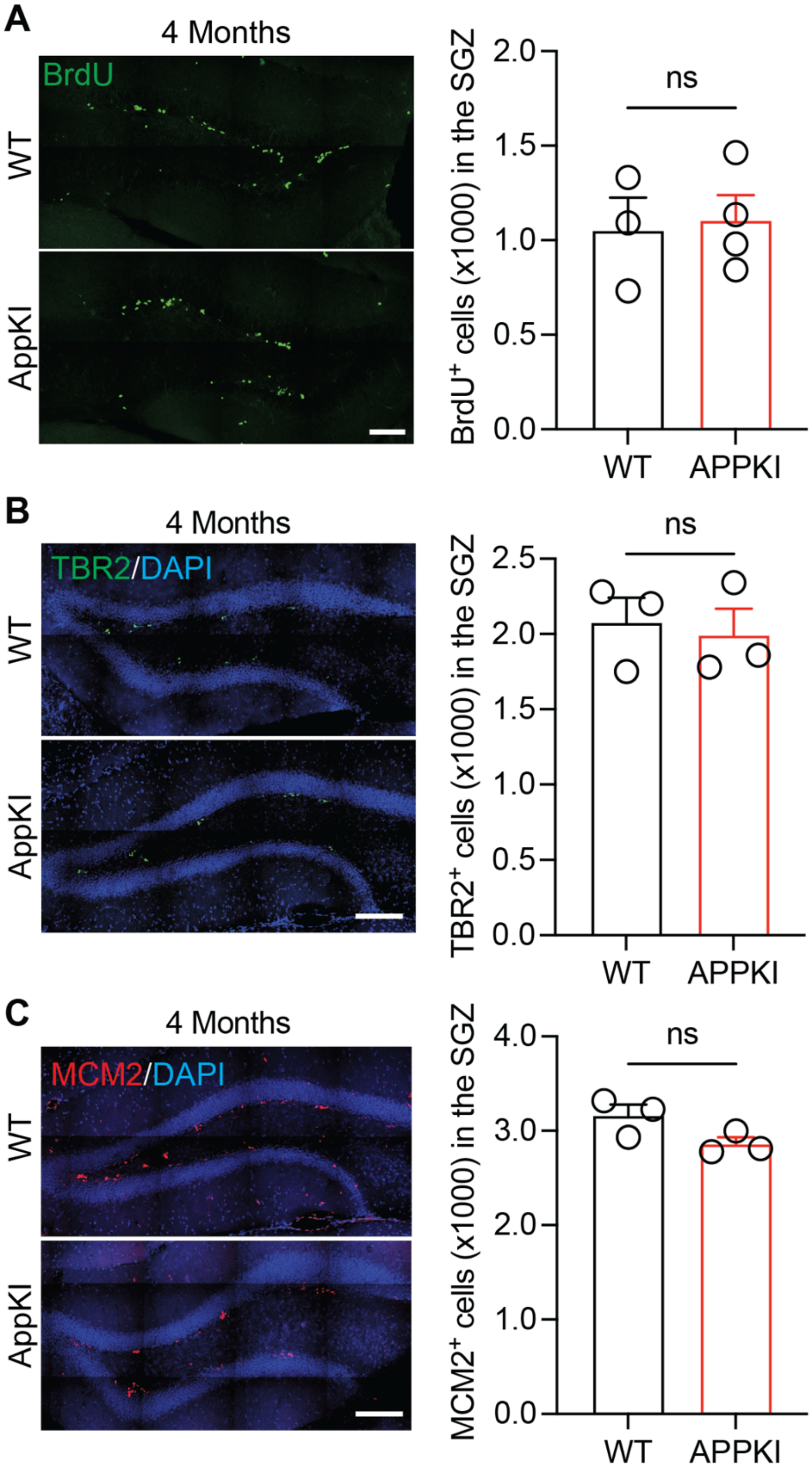


**Supplementary Figure 4 Analysis of proliferation rate of AHN in 4-months-old WT and AppKI mice**

**A Left:** Representative confocal images of BrdU immunostaining in the DG of 4-month-old WT and AppKI mice for proliferation rate analysis (scale bar: 100 μm). **Right:** Quantification of BrdU+ cells in the SGZ for proliferation rate of AHN in 4-month-old WT and AppKI mice (WT = 3, AppKI = 4; 2-tailed unpaired t-test; ns, $p>0.05$).

**B Left:** Representative confocal images of TBR2 immunostaining in the DG of 4-month-old WT and AppKI mice (scale bar: 200 μm). **Right:** Quantification of $TBR2^+$ cells in the SGZ of 4-month-old WT and AppKI mice (WT = 3, AppKI = 3; 2-tailed unpaired t-test; ns, $p>0.05$).

**C Left:** Representative confocal images of MCM2 immunostaining in the DG of 4-month-old WT and AppKI mice (scale bar: 200 μm). **Right:** Quantification of $MCM2^+$ cells in the SGZ of 4-month-old WT and AppKI mice (WT = 3, AppKI = 3; 2-tailed unpaired t-test; ns, $p>0.05$).
